# Consistent inter-protocol differences in resting-state functional connectomes between normal aging and mild cognitive impairment

**DOI:** 10.1101/019646

**Authors:** Angela Tam, Christian Dansereau, AmanPreet Badhwar, Pierre Orban, Sylvie Belleville, Howard Chertkow, Alain Dagher, Alexandru Hanganu, Oury Monchi, Pedro Rosa-Neto, Amir Shmuel, Seqian Wang, John Breitner, Pierre Bellec, Alzheimer’s Disease Neuroimaging Initiative

## Abstract

Resting-state functional connectivity is a promising biomarker for Alzheimer’s disease. However, previous resting-state functional magnetic resonance imaging studies in Alzheimer’s disease and mild cognitive impairment (MCI) have shown limited reproducibility as they have had small sample sizes and substantial variation in study protocol. We sought to identify functional brain networks and connections that could consistently discriminate normal aging from MCI despite variations in scanner manufacturer, imaging protocol, and diagnostic procedure. We therefore pooled four independent datasets, including 112 healthy controls and 143 patients with MCI, systematically testing multiple brain connections for consistent differences. The largest effects associated with MCI involved the ventromedial and dorsomedial prefrontal cortex, striatum, and middle temporal lobe. Compared with controls, patients with MCI exhibited significantly decreased connectivity within the frontal lobe, between frontal and temporal areas, and between regions of the cortico-striatal-thalamic loop. Despite the heterogeneity of methods among the four datasets, we identified robust MCI-related connectivity changes with small to medium effect sizes and sample size estimates recommending a minimum of 150 to 400 total subjects to achieve adequate statistical power. If our findings can be replicated and associated with other established biomarkers of Alzheimer’s disease (e.g. amyloid and tau quantification), then these functional connections may be promising candidate biomarkers for Alzheimer’s disease.

## 1. Introduction

Resting-state connectivity in functional magnetic resonance imaging (fMRI) captures the spatial coherence of spontaneous fluctuations in blood oxygenation. Resting-state fMRI is a promising technique that may be useful as an early biomarker for Alzheimer’s disease (AD), a neurodegenerative process that develops over decades before patients suffer from dementia. The possibility that disturbed resting-state connectivity may be an early marker for AD is supported by studies of mild cognitive impairment (MCI), a disorder characterized by objective cognitive deficits without dementia, i.e., without impairment in activities of daily living. These studies showed altered functional connectivity in MCI compared with cognitively normal elderly (CN) (Bai et al., 2009; Liang et al., 2012; Sorg et al., 2007; Wu et al., 2014), but they relied on small sample sizes (n∼40) and differed in many aspects of their protocols, e.g. recruitment and image acquisition procedures. If resting-state fMRI is to serve as a useful biomarker of AD, or any pathology, for clinical practice or research, we must determine if changes in functional connectivity differences between groups of subjects are robust to such variation in study protocols. Therefore, we sought to identify brain connections that showed consistent MCI-related changes across multiple independent studies. If such connections exist, they may be used as targets to be examined alongside other established AD biomarkers (e.g. amyloid and tau measures) in order to validate resting-state fMRI’s potential as a biomarker for AD.

Resting-state connectivity studies have consistently found decreased connectivity between nodes within the default mode network (DMN) in patients with AD or MCI compared with CN (Bai et al., 2009; Koch et al., 2012; Liang et al., 2012; Sorg et al., 2007; Zhang et al., 2010). Less consistent are reports of alterations in the executive attentional, frontoparietal, and anterior temporal networks (Agosta et al., 2012; Gour et al., 2011; Liang et al., 2012; Sorg et al., 2007; Wu et al., 2014; Zhang et al., 2010) due to the literature’s bias towards investigating the DMN. Further inconsistencies can be found in some studies have reported increased connectivity between the middle temporal lobe and other DMN areas in MCI (Qi et al., 2010), while others have reported decreased connectivity between these same regions (Bai et al., 2009) and others have reported no significant differences between MCI and CN (Koch et al., 2012).

One obvious explanation for such inconsistency may be these studies’ small sample sizes resulting in low statistical power (Kelly et al., 2012). Beyond this, however, there are other methodological differences that may compromise the comparison of results across independent studies. For example, the criteria for recruiting subjects with MCI, e.g. Petersen (2004), vs. NIA-AA recommendations (Albert et al., 2011) may differ among studies. Different study samples may also reflect different socio-cultural characteristics of recruiting sites, e.g., ethnicity, language, diet, socioeconomic status. The fMRI measurements themselves can also be affected by differences in details of the image acquisition such as scanner make and model (Friedman et al., 2006), sequence parameters such as repetition time, flip angle, or acquisition volume (Friedman and Glover, 2006), experimental design such as eyes-open/eyes-closed (Yan et al., 2009) or experiment duration (Van Dijk et al., 2010), and scanning environment such as sound attenuation measures (Elliott et al., 1999), room temperature (Vanhoutte et al., 2006), or head-motion restraint techniques (Edward et al., 2000).

To identify robust changes in resting-state connectivity between MCI and CN, we implemented a pooled analysis of four independent resting-state fMRI datasets (ADNI2 and three small single-site studies) using a weighted average implemented by Willer et al (2010) for meta-analysis. Rather than relying on a priori target regions or connections, we leveraged the large sample size to perform a systematic search of brain connections affected by MCI, an approach termed a “connectome-wide association study” (Shehzad et al., 2014). In addition, we relied on functionally-defined brain parcellations using an automated clustering procedure and we explored the impact of the number of brain clusters (called resolution) on observed differences (Bellec et al., 2014).

## 2. Methods

### 2.1 Participants

We pooled data from four independent studies: the Alzheimer’s Disease Neuroimaging Initiative 2 (ADNI2) sample, two samples from the Centre de recherche de l’institut universitaire de gériatrie de Montréal (CRIUGMa and CRIUGMb), and a sample from the Montreal Neurological Institute (MNI) (Wu et al., 2014). All participants gave their written informed consent to engage in these studies, which were approved by the research ethics board of the respective institutions, and included consent for data sharing with collaborators as well as secondary analysis. Ethical approval was also obtained at the site of secondary analysis (CRIUGM).

The ADNI2 data used in the preparation of this article were obtained from the Alzheimer’s Disease Neuroimaging Initiative (ADNI) database (adni.loni.usc.edu). ADNI was launched in 2003 by the National Institute on Aging, the National Institute of Biomedical Imaging and Bioengineering, the Food and Drug Administration, private pharmaceutical companies and non-profit organizations, as a $60 million, 5-year public-private partnership representing efforts of co-investigators from numerous academic institutions and private corporations. ADNI was followed by ADNI-GO and ADNI-2 that included newer techniques. Subjects included in this study were recruited by ADNI-2 from all 13 sites that acquired resting-state fMRI on Philips scanners across North America. For up-to-date information, see www.adni-info.org.

The combined sample included 112 CN and 143 MCI. In the CN group, the mean age was 72.0 (s.d. 7.0) years, and 38.4% were men. Mean age of the MCI subjects was 72.7 (s.d. 7.7) years, and 50.3% were men. An independent samples t-test did not reveal any significant difference in age between the groups (*t* = 0.759, *p* = 0.448). A chi-squared test revealed a trend towards a significant difference in gender distribution between the groups (*χ*^2^ = 3.627, *p* = 0.057). Note that both age and gender were entered as confounding variables in the statistical analysis below. See Table 1 for sample size and demographic information from the individual studies after passing quality control (for information about the original cohorts before quality control, see Supplementary Table 1).

**Table 1.** Demographic information in all studies after quality control

|  |  | **ADNI2** | **CRIUGMa** | **CRIUGMb** | **MNI** | **Combined sample** |
| --- | --- | --- | --- | --- | --- | --- |
| **CN** | **N** | 49 | 18 | 17 | 15 | 99 |
|  | Mean age (s.d.) | 74.4 (6.8) | 71.2 (8.0) | 70.4 (4.6) | 67.0 (5.7) | 72.0 (7.0) |
|  | Number male (%) | 21 (43%) | 7 (39%) | 2 (12%) | 7 (47%) | 37 (37%) |
|  | MMSE mean (range) | 28.7 (25-30) | 28.8 (27-30) | n/a | 29.1 (n/a) | n/a |
|  | MoCA mean (range) | n/a | 27.8 (22-30) | 28.4 (26-30) | n/a | n/a |
| **MCI** | **N** | 82 | 8 | 21 | 18 | 129 |
|  | Mean age (s.d.) | 71.2 (7.3) | 79.9 (6.1) | 74.8 (7.0) | 71.2 (8.1) | 72.3 (7.6) |
|  | Number male (%) | 43 (52%) | 3 (38%) | 12 (57%) | 7 (39%) | 65 (50%) |
|  | MMSE mean (range) | 28.1 (24-30) | 26.1 (22-29) | n/a | 26.2 (n/a) | n/a |
|  | MoCA mean (range) | n/a | 23.3 (20-29) | 24.8 (16-29) | n/a | n/a |

MMSE: Mini-mental state examination; MoCA: Montreal Cognitive Assessment

All subjects underwent cognitive testing (e.g. memory, language, and executive function) (see Supplementary Table 2 for a list of specific tests used in each study). Exclusion criteria common to all studies included: Contraindications to MRI, presence or history of axis I psychiatric disorders (e.g. depression, bipolar disorder, schizophrenia), presence or history of neurologic disease with potential impact on cognition (e.g. Parkinson’s disease), and presence or history of substance abuse. CN subjects could not meet criteria for MCI or dementia. Those with MCI had memory complaints, objective cognitive loss (based on neuropsychological testing), but had intact functional abilities and did not meet criteria for dementia. In ADNI2, the diagnosis of MCI was made based on an education adjusted abnormal score on the Logical Memory II subscale (Delayed Paragraph Recall, Paragraph A only) from the Wechsler Memory Scale and a Clinical Dementia Rating (CDR) of 0.5. In both CRIUGMa and CRIUGMb, the diagnosis of MCI was made based on scores equal to or greater than 1.5 standard deviations below the mean adjusted for age and education on memory tests. At the MNI, the diagnosis of MCI relied on the Petersen criteria (2004). At both CRIUGMb and MNI, MCI diagnoses were made with input from a neurologist. See the Supplementary Methods for greater details for each study.

### 2.2 Imaging data acquisition

All resting-state fMRI and structural scans were acquired on 3T scanners. We performed analyses on the first usable scan (typically the baseline scan) from ADNI2 and applied clinical diagnoses from the same study time point as the first usable scan for each participant in that dataset. See Table 2 for acquisition parameters for each sample.

**Table 2.**
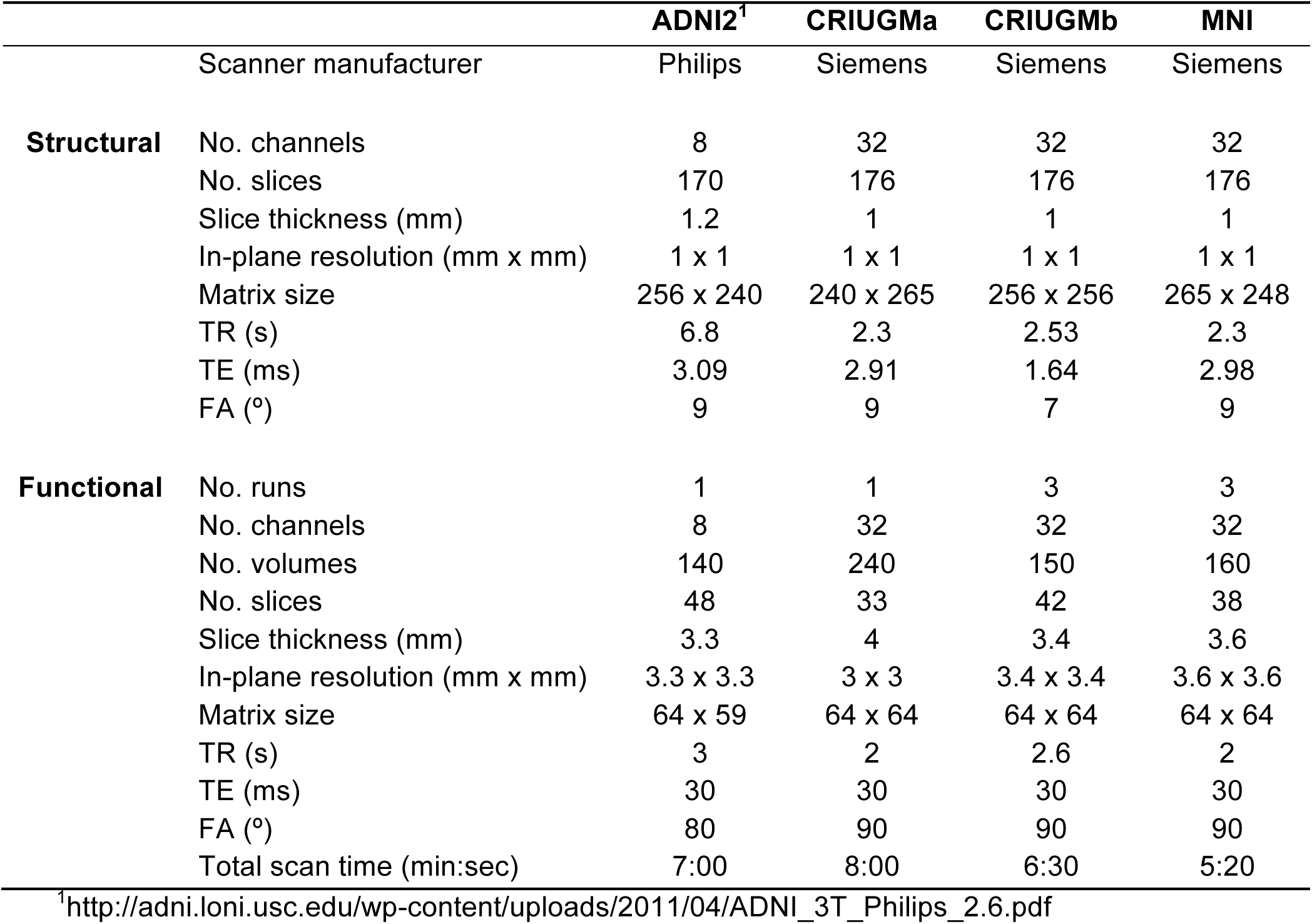
Structural and functional scan acquisition parameters

|  |  | ADNI2<sup>1</sup> | CRIUGMa | CRIUGMb | MNI |
| --- | --- | --- | --- | --- | --- |
|  | Scanner manufacturer | Philips | Siemens | Siemens | Siemens |
| **Structural** | No. channels | 8 | 32 | 32 | 32 |
|  | No. slices | 170 | 176 | 176 | 176 |
|  | Slice thickness (mm) | 1.2 | 1 | 1 | 1 |
|  | In-plane resolution (mm x mm) | 1 x 1 | 1 x 1 | 1 x 1 | 1 x 1 |
|  | Matrix size | 256 x 240 | 240 x 265 | 256 x 256 | 265 x 248 |
|  | TR (s) | 6.8 | 2.3 | 2.53 | 2.3 |
|  | TE (ms) | 3.09 | 2.91 | 1.64 | 2.98 |
|  | FA (°) | 9 | 9 | 7 | 9 |
| **Functional** | No. runs | 1 | 1 | 3 | 3 |
|  | No. channels | 8 | 32 | 32 | 32 |
|  | No. volumes | 140 | 240 | 150 | 160 |
|  | No. slices | 48 | 33 | 42 | 38 |
|  | Slice thickness (mm) | 3.3 | 4 | 3.4 | 3.6 |
|  | In-plane resolution (mm x mm) | 3.3 x 3.3 | 3 x 3 | 3.4 x 3.4 | 3.6 x 3.6 |
|  | Matrix size | 64 x 59 | 64 x 64 | 64 x 64 | 64 x 64 |
|  | TR (s) | 3 | 2 | 2.6 | 2 |
|  | TE (ms) | 30 | 30 | 30 | 30 |
|  | FA (°) | 80 | 90 | 90 | 90 |
|  | Total scan time (min:sec) | 7:00 | 8:00 | 6:30 | 5:20 |

<sup>1</sup>http://adni.loni.usc.edu/wp-content/uploads/2011/04/ADNI_3T_Philips_2.6.pdf

### 2.3 Computational environment

All experiments were performed using the NeuroImaging Analysis Kit (NIAK1) (Bellec et al., 2011) version 0.12.18, under CentOS version 6.3 with Octave2 version 3.8.1 and the Minc toolkit3 version 0.3.18. Analyses were executed in parallel on the “Guillimin” supercomputer4, using the pipeline system for Octave and Matlab (Bellec et al., 2012), version 1.0.2. The scripts used for processing can be found on Github5.

### 2.4 Pre-processing

Each fMRI dataset was corrected for slice timing; a rigid-body motion was then estimated for each time frame, both within and between runs, as well as between one fMRI run and the T1 scan for each subject (Collins and Evans, 1997). The T1 scan was itself non-linearly co-registered to the Montreal Neurological Institute (MNI) ICBM152 stereotaxic symmetric template (Fonov et al., 2011), using the CIVET pipeline (Ad-Dab’bagh et al., 2006). The rigid-body, fMRI-to-T1 and T1-to-stereotaxic transformations were all combined to resample the fMRI in MNI space at a 3 mm isotropic resolution. To minimize artifacts due to excessive motion, all time frames showing a displacement greater than 0.5 mm were removed (Power et al., 2012). A minimum of 50 unscrubbed volumes per run was required for further analysis (13 CN and 14 MCI were rejected). The following nuisance covariates were regressed out from fMRI time series: slow time drifts (basis of discrete cosines with a 0.01 Hz high-pass cut-off), average signals in conservative masks of the white matter and the lateral ventricles as well as the first 3 to 10 principal components (median numbers for ADNI2, CRIUGMa, CRIUGMb, and MNI were 9, 6, 7, and 7 respectively, and accounting for 95% variance) of the six rigid-body motion parameters and their squares (Giove et al., 2009; Lund et al., 2006). The fMRI volumes were finally spatially smoothed with a 6 mm isotropic Gaussian blurring kernel. A more detailed description of the pipeline can be found on the NIAK website6 and Github7.

### 2.5 Bootstrap Analysis of Stable Clusters (BASC)

We applied a BASC to identify clusters that consistently exhibited similar spontaneous BOLD fluctuations in individual subjects, and were spatially stable across subjects. We first applied a region-growing algorithm to reduce each fMRI dataset into a time x space array, with 957 regions (Bellec et al., 2006). BASC replicates a hierarchical Ward clustering 1000 times and computes the probability that a pair of regions fall in the same cluster, a measure called stability. The region × region stability matrix is fed into a clustering procedure to derive consensus clusters, which are composed of regions with a high average probability of being assigned to the same cluster across all replications. At the individual level, the clustering was applied to the similarity of regional time series, which was replicated using a circular block bootstrap. Consensus clustering was applied to the average individual stability matrix to identify group clusters. The group clustering was replicated via bootstrapping of subjects in the group. A consensus clustering was finally applied on the group stability matrix to generate group consensus clusters.

The cluster procedure was carried out at a specific number of clusters (called resolution). Using a “multiscale stepwise selection” (MSTEPS) method (Bellec, 2013), we determined a subset of resolutions that provided an accurate summary of the group stability matrices generated over a fine grid of resolutions: 4, 6, 12, 22, 33, 65, 111 and 208.

### 2.6 Derivation of functional connectomes

For each resolution *K*, and each pair of distinct clusters, the between-clusters connectivity was measured by the Fisher transform of the Pearson’s correlation between the average time series of the clusters. The within-cluster connectivity was the Fisher transform of the average correlation between time series inside the cluster. An individual connectome was thus a *K × K* matrix. See Figure 1a-b for an illustration of a parcellation and associated connectome.

**Figure 1.**
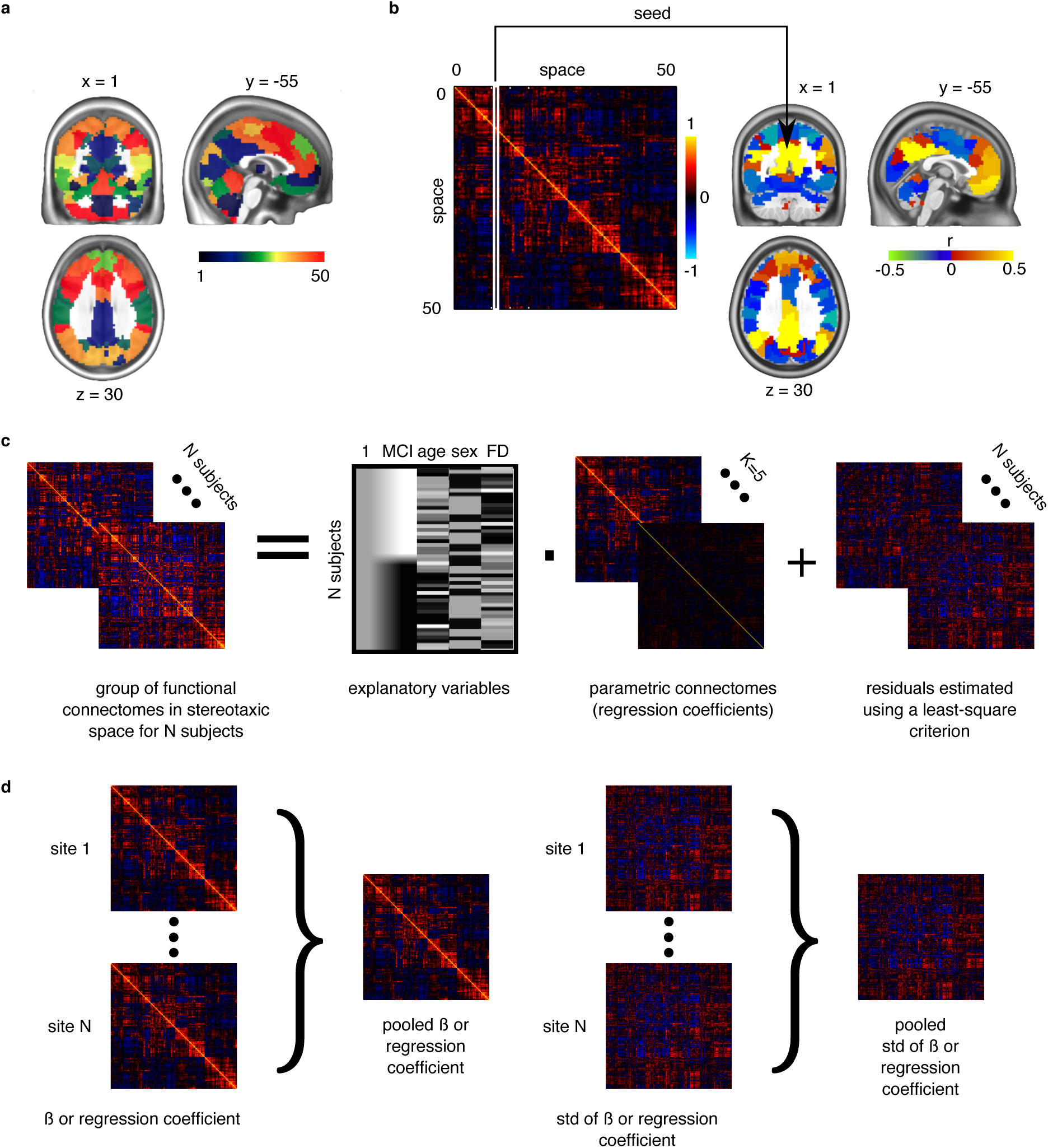
Application of general linear models to connectomes. a) The brain is functionally parcellated into *K* (e.g. 50) clusters generated through a clustering algorithm. b) The connectome is a *K × K* matrix measuring functional connectivity between and within clusters. c) A general linear model is used to test the association between phenotypes and connectomes, independently at each connection, at the group level. d) In a multisite situation, independent site-specific effects are estimated and then pooled through weighted averaging (Willer et al., 2010).

### 2.7 Statistical testing

To test for differences between MCI and CN at a given resolution, we used a general linear model (GLM) for each connection between two clusters. The GLM included an intercept, the age and sex of participants, and the average frame displacement of the runs involved in the analysis. The contrast of interest (MCI – CN) was represented by a dummy covariate coding the difference in average connectivity between the two groups. All covariates except the intercept were corrected to a zero mean (Figure 1c). We estimated the effects independently at each site and pooled them through inverse variance based weighted averaging (Willer et al., 2010) (Figure 1d). Due to the harmonization of protocols across the 13 ADNI2 sites, all ADNI2 sites were combined to form one single site.

Resolutions containing fewer than 50 clusters have been suggested to have higher sensitivity based on prior independent work (Bellec et al., 2014). The GLM was first applied at an a priori resolution of *K* = 33, which was the lowest number of clusters for which the default mode network could be clearly decomposed into subnetworks (Supplementary Figure S1, visit Figshare for 3D volumes of brain parcellations8 and see Supplementary Table 3 for a list of the 33 clusters and their numerical IDs). The false-discovery rate (FDR) across connections was controlled at *q*^*FDR*^ ≤ 0.1 (Benjamini and Hochberg, 1995). In addition to the analysis at resolution 33, we assessed the impact of that parameter by replicating the GLM analysis at the 7 resolutions selected by MSTEPS (Supplementary Figure S2). We implemented an omnibus test (α ≤ 0.05) to assess the overall presence of significant differences between groups, pooling FDR results across all resolutions (Bellec et al., 2014).

## 3. Results

### 3.1 Functional connectivity differences between MCI and CN

The omnibus test pooling significant differences in connectivity between MCI and CN across all resolutions was significant at α ≤ 0.05 (*p* = 0.013). In line with prior observations on independent datasets (Bellec et al., 2014), resolutions containing fewer than 50 clusters were associated with a higher rate of discovery (Figure 2). At resolution 33, significant group differences between MCI and CN were seen across the whole brain, save for the occipital lobe and cerebellum (Figure 3a). Four brain clusters were associated with 64% of all significant changes found across the connectome: the ventromedial prefrontal cortex, striatum, dorsomedial prefrontal cortex, and middle temporal lobe (Figures 3b, 3c, Supplementary Table 3). Supplementary Table 4 contains a list of parcels that account for all non-redundant significant connectivity differences between MCI and CN. For example, the first-ranked seed (ventromedial prefrontal cortex) was associated with 20% of connections that differ between the groups. The second-ranked seed (striatum) was associated with an additional 17.8% of connectivity differences that did not overlap with or were not previously accounted for by the first seed. Note that if a given parcel was associated with a significant effect with another region that ranked in the table, then that parcel may not be listed in the table (i.e. this table is not a comprehensive list of parcels that show significant effects, as a given parcel may involve a region in the table at a higher rank which already accounted for its effects). Given that the top four clusters explained the majority of the findings, they were further characterized in seed-based connectivity analyses, which revealed that MCI showed decreased connectivity between dorsal and ventral frontal areas, between frontal and temporal regions, and between areas of the cortico-striatal-thalamic loop (Figure 4). More specifically, in MCI compared to CN, the ventromedial prefrontal cortex displayed significantly reduced connectivity with dorsal frontal regions and the sensorimotor cortex (Figure 4a). The striatum in MCI exhibited decreased connectivity with the sensorimotor cortex, thalamus, and frontal and parietal regions (Figure 4b). MCI also showed reduced connectivity between the dorsomedial prefrontal cortex with temporal lobe regions, ventral frontal areas, and the cuneus (Figure 4c). Lastly, in MCI, the middle temporal lobe displayed significantly decreased connectivity with the posterior cingulate, precuneus, hippocampus, and frontal areas (Figure 4d).

**Figure 2.**
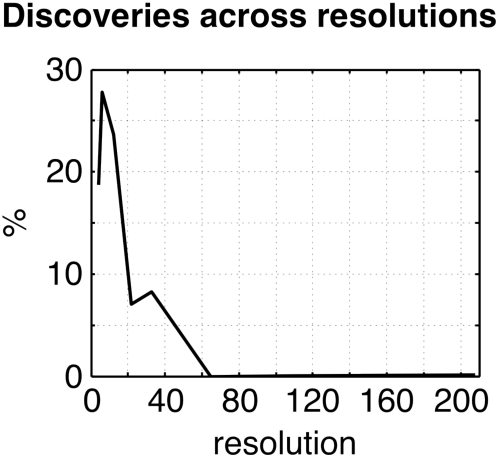
Plot of the percentage of connections identified as significant by the statistical comparison between MCI and CN across the connectome (*q*^*FDR*^ ≤ 0.1), as a function of the resolutions selected by MSTEPS.

**Figure 3.**
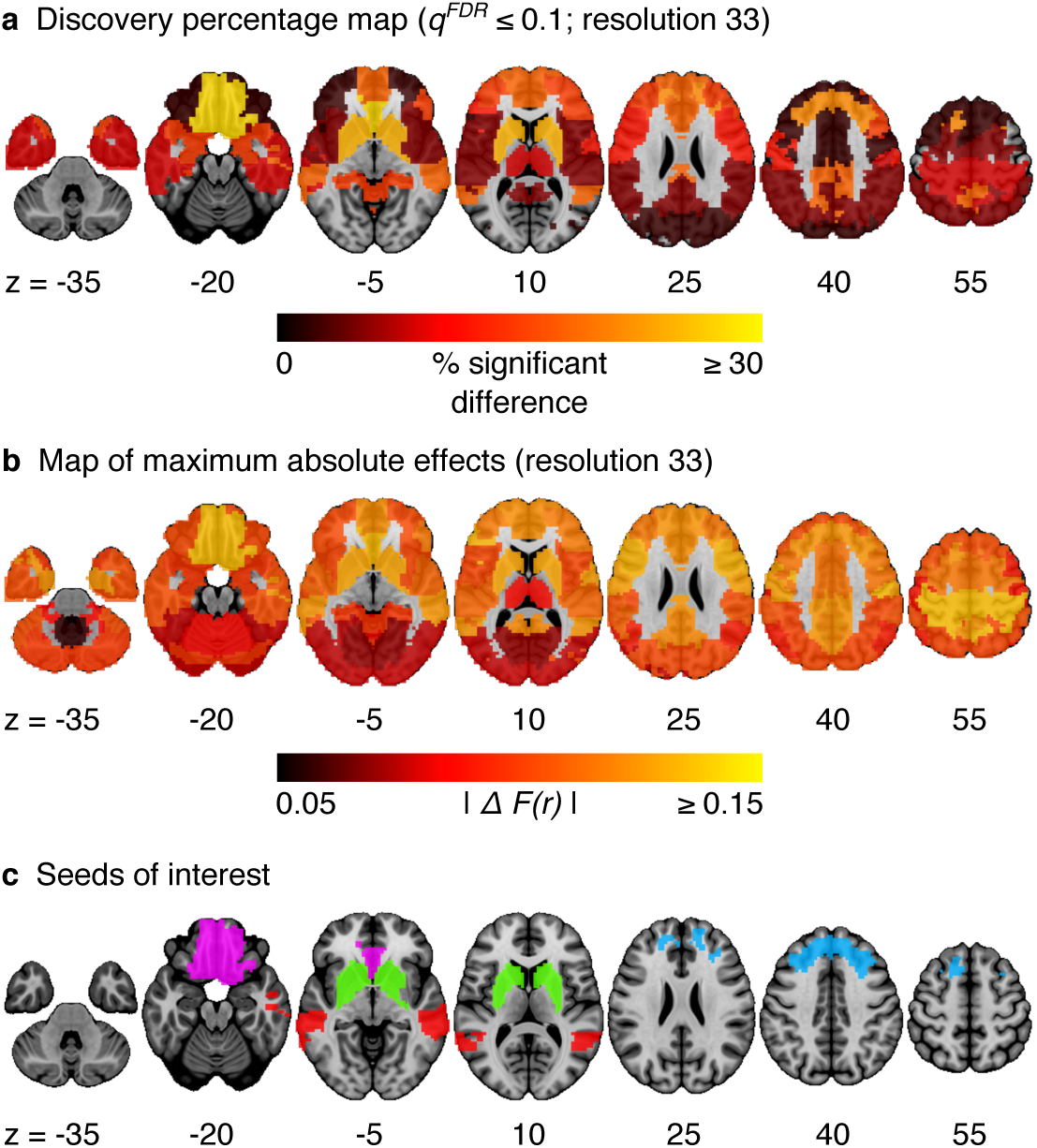
a) Map of the percentage of connections associated with a given cluster and identified as significant by the statistical comparison between MCI and CN, at a resolution of 33 clusters (*q*^*FDR*^ ≤ 0.1). ΔF(r) signifies the difference in Fisher-transformed correlation values between the groups. b) Maximum absolute difference in average connectivity between MCI and CN, across all connections associated with a cluster, at resolution 33. c) Four clusters of interest (ventromedial prefrontal cortex, striatum, dorsomedial prefrontal cortex, middle temporal lobe) selected out of 33 for further characterization.

**Figure 4.**
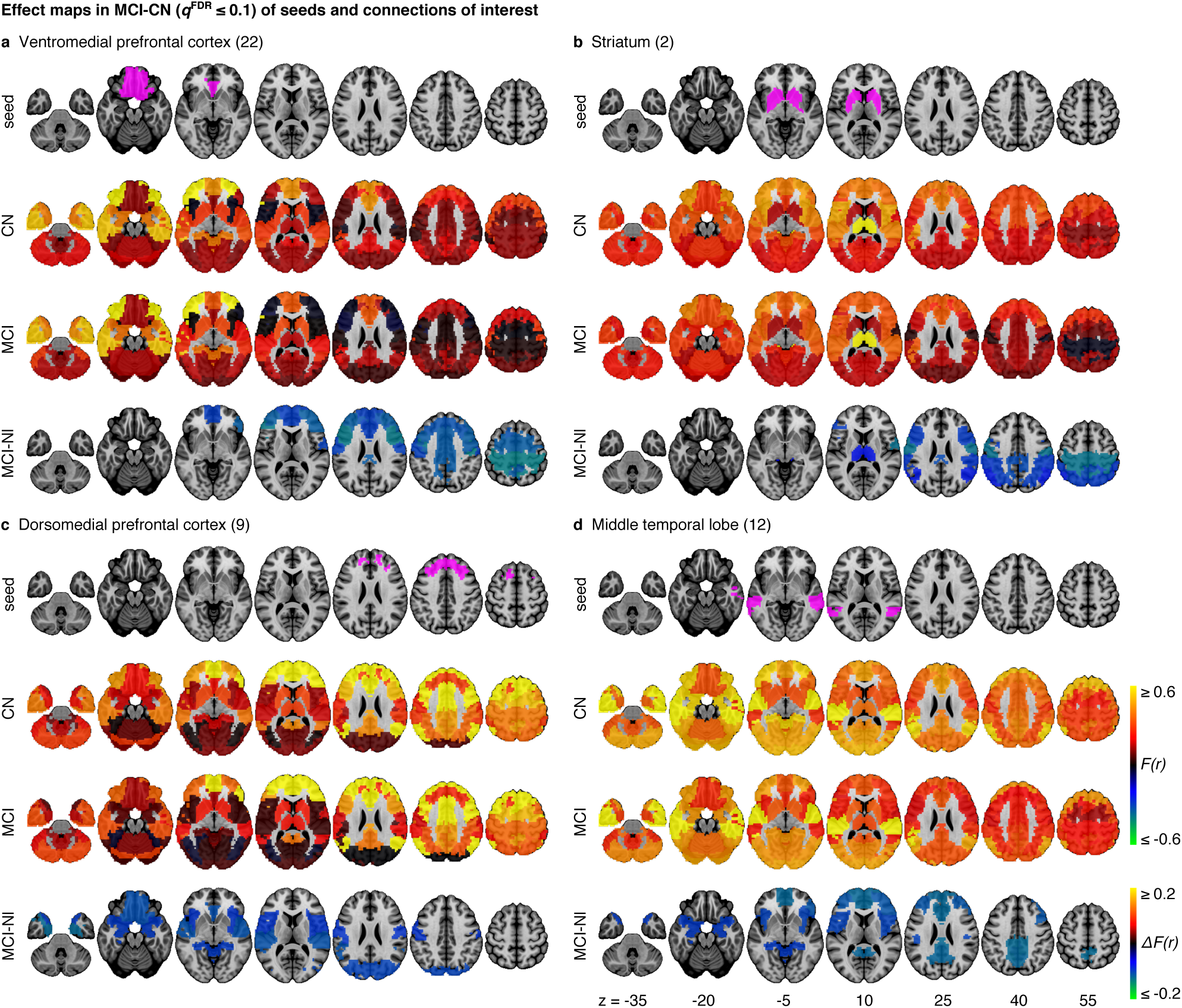
Effect maps for a selection of four seeds that show effects related to MCI at resolution 33. Effect maps reveal the spatial distribution of the changes in functional connectivity for a) the ventromedial prefrontal cortex, b) striatum, c) the dorsomedial prefrontal cortex, and d) the middle temporal lobe. All connections shown in the maps of difference in average connectivity between MCI and CN are significant at *q*^*FDR*^ ≤ 0.1. For each panel, the top line maps the spatial location of the seed region in magenta, the second and third lines show the connectivity (Fisher-transformed correlation values, *F(r)*) between the designated seed region and the rest of the brain in CN and MCI respectively, and the fourth line shows a difference map between MCI and CN (difference in Fisher-transformed correlation values, *ΔF(r)*). The numbers in parentheses refer to the numerical IDs of the clusters in the 3D parcellation volume, as listed in Supplementary Table 3.

### 3.2 Sample-specific effects

The statistical model we used to combine GLM analyses across sites was based on a (weighted) average. The possibility thus existed that an effect would be significant in the pooled analysis because it was driven by a very strong effect in a single sample, instead of being consistent across all samples. By examining effects in each sample independently, we found that MCI-related connectivity changes that surpassed the FDR threshold in the pooled analysis showed similar trends in all samples across seeds and connections, where the independent MCI samples consistently exhibited decreased connectivity compared to the CN samples (Figure 5, Supplementary Figures S3,S4,S5,S6). For example, the pooled analysis revealed that, compared to CN, MCI exhibited significantly reduced connectivity between the ventromedial prefrontal cortex (the region in which connectivity was most affected by MCI) and dorsal frontal regions, including the sensorimotor cortex. This change appeared to be consistent in all four independent samples (Supplementary Figure S3). For this particular seed, the change in connectivity was mainly due to regions with positive correlations in CN having smaller correlation values closer to zero in MCI in the individual samples (Supplementary Figure S3).

**Figure 5.**
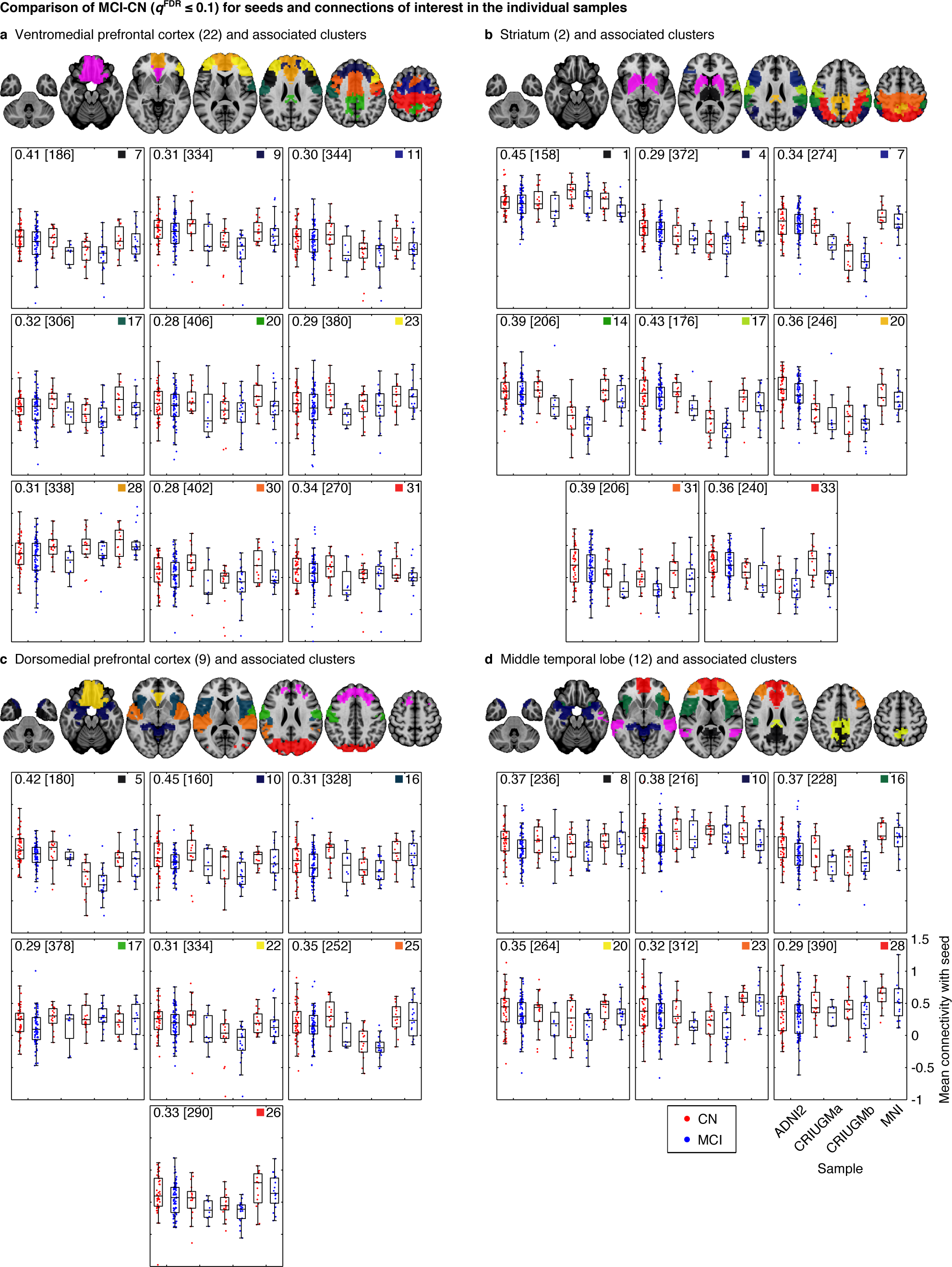
Mean connectivity of the a) ventromedial prefrontal cortex, b) striatum, c) dorsomedial prefrontal cortex, and d) middle temporal lobe with their associated connections in CN and MCI in the independent samples. Each map displays the seed in pink and the clusters (in other colors) whose connectivity with the seed significantly differed between CN and MCI in the pooled analysis. The box-whisker plots display the mean connectivity (Fisher-transformed correlation values) between the seed and a significant parcel, overlaid over individual data points, in the CN and MCI groups in the ADNI2, CRIUGMa, CRIUGMb, and MNI samples. We also report the Cohen’s *d* (a weighted average of the effect sizes per sample) followed by a sample size estimate (for 80% power, balanced groups, bilateral tests, Gaussian distributions, and α = 0.05) in square brackets in the top-left corner of each plot. The numbers in parentheses in the titles refer to the numerical IDs of the seeds in the 3D parcellation volume, as listed in Supplementary Table 3. Similarly, each plot is labeled with a number in the top-right corner, corresponding to the number assigned to the cluster in that table, and a colored square corresponding to the parcel of the same color from the map.

### 3.3 Effect sizes and sample size estimates

We measured the effect sizes of the difference between groups at each significant connection by calculating Cohen’s *d*, via a weighted average of the effect sizes per individual sample. We found small to medium effect sizes, ranging from *d* = 0.28 to *d* = 0.45, with an average effect size of *d* = 0.35. Note that these effect sizes are potentially inflated since we have focussed on significant results only. We also calculated the sample sizes required to achieve 80% power, based on the effect sizes estimated by Cohen’s *d*, the assumption of balanced groups, Gaussian distributions, bilateral tests, and α = 0.05, for each connection. We found that the estimated sample sizes ranged from approximately 150 to upwards of 400 total subjects. See Figure 5 for Cohen’s *d* and sample size estimates for each significant connection that was reported in Figure 4.

### 3.4 Effect of resolution on the GLM

The percentage of discoveries in significant differences between MCI and CN across the connectomes varied markedly as a function of resolution, as selected by the MSTEPS procedure. Higher resolutions were associated with fewer discoveries, especially beyond resolution 65 (Supplementary Figure S7a). By contrast, the maximal amplitude of differences in average connectivity associated with a particular cluster did not decrease substantially, and sometimes increased, when the resolution increased. The decrease in percentage of discovery thus likely reflected a cost associated with an increased number of multiple comparisons in the FDR procedure, rather than a loss in signal quality (Supplementary Figure S7b). Regarding the clusters that were selected for our seed-based analyses (the ventromedial prefrontal cortex, striatum, dorsomedial prefrontal cortex, and middle temporal lobe), the associated effect maps (without statistical threshold) were highly consistent across different resolutions (Supplementary Figures S8-S9), with the potential exception of very low resolutions where, for example, a relatively small cluster like the caudate nuclei got merged with a large distributed cortical network. This also replicated a prior study on the effect of multiresolution parcellations on GLM analysis (Bellec et al., 2014).

## 4. Discussion

We report resting-state functional connectivity differences in the ventromedial prefrontal cortex, striatum, dorsomedial prefrontal cortex, and middle temporal lobe between MCI and CN subjects that were consistent across multiple studies. Despite protocol differences, we found that MCI exhibited reduced connectivity within the frontal lobe, between frontal and temporal regions, and among areas of the cortico-striatal-thalamic loop compared to CN. Previous studies suggested these altered patterns of functional connectivity in MCI may result from the coevolution of multiple AD-associated biological processes, namely structural degeneration (Coupé et al., 2012; Pievani et al., 2010), neurofibrillary and amyloid pathologies (Small et al., 2006), and cerebrovascular dysfunction (Villeneuve and Jagust, 2015).

The ventromedial prefrontal cortex showed the greatest connectivity changes with other brain areas. Decreased functional connectivity between the ventromedial prefrontal cortex and dorsal regions could reflect degeneration in gray matter and in white matter tracts connecting these areas. Longitudinal studies have shown greater prefrontal cortex atrophy in MCI over time, as well as in those transitioning to AD, compared to CN (Carmichael et al., 2013; McDonald et al., 2009). Cortico-cortical white matter bundles, e.g. superior longitudinal fasciculus, have also been demonstrated to degenerate in patients with MCI and AD (Pievani et al., 2010). Additionally, functional connectivity changes may reflect the regional effect of increased amyloid burden (Lim et al., 2014; Sheline et al., 2010b), and PIB-PET work has shown the frontal lobe to be one of the first regions in which amyloid accumulates in autosomal dominant AD mutation carriers (Bateman et al., 2012). Our results may also be due to neurofibrillary pathology as it typically appears in the prefrontal cortex during MCI (Bossers et al., 2010). Lastly, cerebral hypoperfusion in the frontal lobe of MCI (Chao et al., 2009) may have contributed to our results.

As expected, a large effect was also observed in the temporal lobe, a region known to suffer from significant AD pathology in preclinical phases (Guillozet et al., 2003). Compared to CN, we found that MCI displayed reduced connectivity between the middle temporal lobe and large parts of the frontal lobe, hippocampus, posterior cingulate and precuneus. This conforms to previous resting-state studies in MCI and AD that have shown effects in the default mode network (Koch et al., 2012; Sorg et al., 2007). Again, we observed functional disconnection between the temporal and frontal lobes in MCI when the dorsomedial prefrontal cortex was selected as a seed. Structural connectivity may explain the functional connectivity changes in these areas, since degeneration of white matter tracts between frontal and temporal regions, e.g. the uncinate fasciculus, occurs with the progression from MCI to AD and correlates with episodic memory impairment in MCI (Pievani et al., 2010; Rémy et al., 2015). Furthermore, examining the integrity of the arcuate fasciculus, a major language tract that connects the frontal and temporal lobes (Dick and Tremblay, 2012), might reveal a biological basis for language impairments such as word-finding difficulties in MCI and AD, (Nutter-Upham et al., 2008). Brain areas that subserve language function could be important targets to investigate given recent evidence that multilingualism, like other forms of cognitive reserve, may help delay the onset of AD (Chertkow et al., 2010).

Unexpectedly, we also found significant effects in the striatum, which showed reduced connectivity in MCI with sensorimotor cortex, frontal and parietal regions, and thalamus. While not initially expected, these findings may reflect earlier observations that regions within the cortico-striatal-thalamic loops are vulnerable to AD pathology. For example, previous work demonstrated the presence of substantial amyloid burden in the striatum in both autosomal dominant and sporadic forms of AD (H. Braak and E. Braak, 1990; Villemagne et al., 2009), and the striatum may be the first region in which amyloid deposition occurs in autosomal dominant AD (Bateman et al., 2012; Klunk et al., 2007). Furthermore, significant neurodegeneration is known to occur with AD in the striatum and thalamus (de Jong et al., 2008; Madsen et al., 2010), so our results might reflect the brain’s capacity for functional plasticity in response to amyloid or neurodegeneration in these regions. Motor cortex hyperexcitability has also been shown in AD, and this suggests that inhibitory circuits leading to the motor cortex may be affected in the disease (Ferreri et al., 2011). Our results may support that theory. Additionally, our findings may represent a biological basis for the cognitive and motor symptoms of MCI (Aggarwal et al., 2006) since the striatum and the rest of the basal ganglia have been implicated in stimulus-response associative learning and memory and motor skill acquisition and execution (Doyon et al., 2009; Packard and Knowlton, 2002). Future research should examine the potential relationship between connectivity in the cortico-striatal-thalamic loops and motor function in MCI and AD.

The consistency of our findings contrasted with previous, smaller single-site studies that have variously reported decreased and increased connectivity. The reports of increased connectivity (Bai et al., 2009; Gour et al., 2011; Qi et al., 2010) may have reflected unique attributes of particular protocols or the choices made with respect to pre-processing steps, for example using global signal regression (Saad et al., 2012).

Among our study’s limitations is its cross-sectional nature, which precludes inference that the functional changes we found would necessarily predict progression towards Alzheimer’s dementia. Furthermore, MCI has many underlying causes aside from AD. It is possible that some subjects in our cohort had cognitive impairments due to Lewy Body dementia, for example. However, all samples in the current study had inclusion criteria that enriched for subjects that had MCI likely due to AD and excluded MCI subjects with other co-morbidities, such as depression or Parkinson’s disease. Also, we did not account for structural atrophy, despite a bias for increased detection in functional differences due to differences in underlying structure (Dukart and Bertolino, 2014). However MCI-related gray matter changes likely co-localize to some extent with functional changes, and the aim of our work was to map out functional changes rather than study their interaction with atrophy. We did not account for other variables, such as APOE genotype (Sheline et al., 2010a), amyloid deposition (Sheline et al., 2010b), presence of neurofibrillary tangles (Maruyama et al., 2013), and cerebrovascular mechanisms (Villeneuve and Jagust, 2015). At least some of these could potentially have explained the observed MCI-related functional connectivity changes as part of an underlying disease mechanism. Large-scale multimodal studies, incorporating genomics, proteomics, and multimodal imaging will be needed to identify the interactions between these and other physiological facets of the pathology. Despite combining several samples together, we still only achieved relatively limited power, given that sample size estimates required at least 150 to 400 total subjects to consistently identify effects between groups. Lastly, because of the explorative approach used in our study, the resulting estimates of effect sizes may have been inflated and discussion of possible pathological mechanisms for our findings was speculative. However, our discoveries may be used as follow-up targets in future work. Upcoming research should not only attempt to verify our findings by using these regions and their associated connections with hypothesis-driven approaches (e.g. seed-based correlation analyses), but also to extend them to cohorts that include Alzheimer’s dementia and other clinical populations (e.g. CN with significant amyloid deposition) and to longitudinal studies that characterize individuals’ progression to dementia. Finally, future studies should aim to determine whether our findings are associated with established biomarkers of AD (e.g. amyloid and tau quantification) in order to probe the potential of these functional connections as biomarkers.

Overall, our results supported previous findings of DMN connectivity changes in AD and MCI (Greicius et al., 2004; Sorg et al., 2007), given that three of the identified seeds (ventromedial prefrontal cortex, dorsomedial prefrontal cortex, middle temporal lobe) are part of this network. It is noteworthy, however, that our strongest observed effects reported were not in the same DMN regions typically described in earlier resting-state studies of MCI and AD, viz, posterior cingulate/precuneus (Sheline et al., 2010b; Zhang et al., 2010). Unexpected changes were also found in the striatum, and this may reflect the advantages of “mining” the whole-brain connectome to search for new biomarkers of mild cognitive impairment and possibly the early progression of the pathophysiologic substrate of Alzheimer’s disease. If confirmed, our results could suggest the utility of these regions in resting-state fMRI as a biomarker endpoint in clinical trials.

## Acknowledgements

We thank Sven Joubert, Isabelle Rouleau, and Sophie Benoit for their contribution in collection of the CRIUGMa dataset. We thank Tom Nichols for his comments and suggestions. Collection of the CRIUGMb dataset was supported by the Alzheimer’s Society of Canada. Collection of the MNI dataset was supported by Industry Canada/Montreal Neurological Institute Centre of excellence in commercialization and research. This work was supported by CIHR (133359) and NSERC (436141-2013).

Data collection and sharing for this project was funded by the Alzheimer’s Disease Neuroimaging Initiative (ADNI) (National Institutes of Health Grant U01 AG024904) and DOD ADNI (Department of Defense award number W81XWH-12-2-0012). ADNI is funded by the National Institute on Aging, the National Institute of Biomedical Imaging and Bioengineering, and through generous contributions from the following: Alzheimer’s Association; Alzheimer’s Drug Discovery Foundation; Araclon Biotech; BioClinica, Inc.; Biogen Idec Inc.; Bristol-Myers Squibb Company; Eisai Inc.; Elan Pharmaceuticals, Inc.; Eli Lilly and Company; EuroImmun; F. Hoffmann-La Roche Ltd and its affiliated company Genentech, Inc.; Fujirebio; GE Healthcare; IXICO Ltd.; Janssen Alzheimer Immunotherapy Research & Development, LLC.; Johnson & Johnson Pharmaceutical Research & Development LLC.; Medpace, Inc.; Merck & Co., Inc.; Meso Scale Diagnostics, LLC.; NeuroRx Research; Neurotrack Technologies; Novartis Pharmaceuticals Corporation; Pfizer Inc.; Piramal Imaging; Servier; Synarc Inc.; and Takeda Pharmaceutical Company. The Canadian Institutes of Health Research is providing funds to support ADNI clinical sites in Canada. Private sector contributions are facilitated by the Foundation for the National Institutes of Health (<u>www.fnih.org</u>). The grantee organization is the Northern California Institute for Research and Education, and the study is coordinated by the Alzheimer’s Disease Cooperative Study at the University of California, San Diego. ADNI data are disseminated by the Laboratory for Neuro Imaging at the University of Southern California.

## Footnotes

1 <u>http://simexp.github.io/niak/</u>

2 <u>http://gnu.octave.org</u>

3 <u>http://www.bic.mni.mcgill.ca/ServicesSoftware/ServicesSoftwareMincToolKit</u>

4 <u>http://www.calculquebec.ca/en/resources/compute-servers/guillimin</u>

5 <u>https://github.com/SIMEXP/mcinet</u>

6 <u>http://niak.simexp-lab.org/pipe_preprocessing.html</u>

7 <u>https://github.com/SIMEXP/mcinet/tree/master/preprocess</u>

8 <u>http://dx.doi.org/10.6084/m9.figshare.1480461</u>

## References

Ad-Dab’bagh, Y., Einarson, D., Lyttelton, O., Muehlboeck, J.S., Mok, K., Ivanov, O., Vincent, R.D., Lepage, C., Lerch, J., Fombonne, E., Evans, A.C., 2006. The CIVET image-processing environment: a fully automated comprehensive pipeline for anatomical neuroimaging research. Proceedings of the th annual meeting of the organization for human brain mapping 2266.

Aggarwal, N.T., Wilson, R.S., Beck, T.L., Bienias, J.L., Bennett, D.A., 2006. Motor dysfunction in mild cognitive impairment and the risk of incident Alzheimer disease. Arch Neurol 63, 1763–1769. doi:10.1001/archneur.63.12.1763

Agosta, F., Pievani, M., Geroldi, C., Copetti, M., Frisoni, G.B., Filippi, M., 2012. Resting state fMRI in Alzheimer’s disease: beyond the default mode network. Neurobiology of Aging 33, 1564–1578. doi:10.1016/j.neurobiolaging.2011.06.007

Albert, M.S., DeKosky, S.T., Dickson, D., Dubois, B., Feldman, H.H., Fox, N.C., Gamst, A., Holtzman, D.M., Jagust, W.J., Petersen, R.C., Snyder, P.J., Carrillo, M.C., Thies, B., Phelps, C.H., 2011. The diagnosis of mild cognitive impairment due to Alzheimer’s disease: recommendations from the National Institute on Aging-Alzheimer’s Association workgroups on diagnostic guidelines for Alzheimer’s disease. Alzheimers Dement 7, 270–279. doi:10.1016/j.jalz.2011.03.008

Bai, F., Watson, D.R., Yu, H., Shi, Y., Yuan, Y., Zhang, Z., 2009. Abnormal resting-state functional connectivity of posterior cingulate cortex in amnestic type mild cognitive impairment. Brain Research 1302, 167–174. doi:10.1016/j.brainres.2009.09.028

Bateman, R.J., Xiong, C., Benzinger, T.L.S., Fagan, A., Goate, A., Fox, N.C., Marcus, D.S., Cairns, N.J., Xie, X., Blazey, T.M., Holtzman, D.M., Santacruz, A., Buckles, V., Oliver, A., Moulder, K.,Aisen, P.S., Ghetti, B., Klunk, W.E., McDade, E., Martins, R.N., Masters, C.L., Mayeux, R., Ringman, J.M., Rossor, M.N., Schofield, P.R., Sperling, R.A., Salloway, S., Morris, J.C., 2012. Clinical and Biomarker Changes in Dominantly Inherited Alzheimer’s Disease. N. Engl. J. Med. 367, 795–804. doi:10.1056/NEJMoa1202753

Bellec, P., 2013. Mining the Hierarchy of Resting-State Brain Networks: Selection of Representative Clusters in a Multiscale Structure. Pattern Recognition in Neuroimaging 54–57.

Bellec, P., Benhajali, Y., Carbonell, F., Dansereau, C., Albouy, G., Pelland, M., Craddock, C., Collignon, O., Doyon, J., Stip, E., Orban, P., 2014. Multiscale statistical testing for connectome-wide association studies in fMRI. arXiv preprint arXiv:1409.2080.

Bellec, P., Carbonell, F.M., Perlbarg, V., Lepage, C., Lyttelton, O., Fonov, V., Janke, A., Tohka, J., Evans, A.C., 2011. A neuroimaging analysis kit for Matlab and Octave. Proceedings of the th International Conference on Functional Mapping of the Human Brain.

Bellec, P., Lavoie-Courchesne, S., Dickinson, P., Lerch, J.P., Zijdenbos, A.P., Evans, A., 2012. The pipeline system for Octave and Matlab (PSOM): a lightweight scripting framework and execution engine for scientific workflows. Front Neuroinform 6, 7. doi:10.3389/fninf.2012.00007

Bellec, P., Perlbarg, V., Jbabdi, S., Pélégrini-Issac, M., Anton, J.-L., Doyon, J., Benali, H., 2006. Identification of large-scale networks in the brain using fMRI. NeuroImage 29, 1231–1243. doi:10.1016/j.neuroimage.2005.08.044

Benjamini, Y., Hochberg, Y., 1995. Controlling the false discovery rate: a practical and powerful approach to multiple testing. Journal of the Royal Statistical Society. Series B (Methodological) 289–300.

Bossers, K., Wirz, K.T.S., Meerhoff, G.F., Essing, A.H.W., van Dongen, J.W., Houba, P., Kruse, C.G., Verhaagen, J., Swaab, D.F., 2010. Concerted changes in transcripts in the prefrontal cortex precede neuropathology in Alzheimer’s disease. Brain 133, 3699–3723. doi:10.1093/brain/awq258

Braak, H., Braak, E., 1990. Alzheimer’s Disease: Striatal Amyloid Deposits and Neurofibrillary Changes. Journal of Neuropathology & Experimental Neurology 49, 215.

Carmichael, O., McLaren, D.G., Tommet, D., Mungas, D., Jones, R.N., .for the Alzheimer’s Disease Neuroimaging Initiative, 2013. Coevolution of brain structures in amnestic mild cognitive impairment. NeuroImage 66, 449–456. doi:10.1016/j.neuroimage.2012.10.029

Chao, L.L., Pa, J., Duarte, A., Schuff, N., Weiner, M.W., Kramer, J.H., Miller, B.L., Freeman, K.M., Johnson, J.K., 2009. Patterns of cerebral hypoperfusion in amnestic and dysexecutive MCI. Alzheimer Disease & Associated Disorders 23, 245–252. doi:10.1097/WAD.0b013e318199ff46

Chertkow, H., Whitehead, V., Phillips, N., Wolfson, C., Atherton, J., Bergman, H., 2010.

Multilingualism (But Not Always Bilingualism) Delays the Onset of Alzheimer Disease: Evidence From a Bilingual Community. Alzheimer Disease & Associated Disorders 24, 118–125. doi:10.1097/WAD.0b013e3181ca1221

Collins, D.L., Evans, A.C., 1997. Animal: Validation and Applications of Nonlinear Registration-Based Segmentation. Int. J. Patt. Recogn. Artif. Intell. 11, 1271–1294. doi:10.1142/S0218001497000597

Coupé, P., Eskildsen, S.F., Manjón, J.V., Fonov, V.S., Collins, D.L., for the Alzheimer’s Disease Neuroimaging Initiative, 2012. Simultaneous segmentation and grading of anatomical structures for patient“s classification: application to Alzheimer”s disease. NeuroImage 59, 3736–3747. doi:10.1016/j.neuroimage.2011.10.080

de Jong, L.W., van der Hiele, K., Veer, I.M., Houwing, J.J., Westendorp, R.G.J., Bollen, E.L.E.M., de Bruin, P.W., Middelkoop, H.A.M., van Buchem, M.A., van der Grond, J., 2008. Strongly reduced volumes of putamen and thalamus in Alzheimer’s disease: an MRI study. Brain 131, 3277–3285. doi:10.1093/brain/awn278

Dick, A.S., Tremblay, P., 2012. Beyond the arcuate fasciculus: consensus and controversy in the connectional anatomy of language. Brain 135, 3529–3550. doi:10.1093/brain/aws222

Doyon, J., Bellec, P., Amsel, R., Penhune, V., Monchi, O., Carrier, J., Lehéricy, S., Benali, H., 2009. Contributions of the basal ganglia and functionally related brain structures to motor learning. Behavioural Brain Research 199, 61–75. doi:10.1016/j.bbr.2008.11.012

Dukart, J., Bertolino, A., 2014. When Structure Affects Function – The Need for Partial Volume Effect Correction in Functional and Resting State Magnetic Resonance Imaging Studies. PLoS ONE 9, e114227-18. doi:10.1371/journal.pone.0114227

Edward, V., Windischberger, C., Cunnington, R., Erdler, M., Lanzenberger, R., Mayer, D., Endl, W., Beisteiner, R., 2000. Quantification of fMRI artifact reduction by a novel plaster cast head holder. Hum. Brain Mapp. 11, 207–213.

Elliott, M.R., Bowtell, R.W., Morris, P.G., 1999. The effect of scanner sound in visual, motor, and auditory functional MRI. Magn Reson Med 41, 1230–1235. doi:10.1002/(SICI)1522-2594(199906)41:6<1230::AID-MRM20>3.0.CO;2-1

Ferreri, F., Pasqualetti, P., Määttä, S., Ponzo, D., Guerra, A., Bressi, F., Chiovenda, P., Del Duca, M., Giambattistelli, F., Ursini, F., Tombini, M., Vernieri, F., Rossini, P.M., 2011. Motor cortex excitability in Alzheimer’s disease: a transcranial magnetic stimulation follow-up study. Neuroscience Letters 492, 94–98. doi:10.1016/j.neulet.2011.01.064

Fonov, V., Evans, A., Botteron, K., Almli, C.R., McKinstry, R.C., Collins, D.L., 2011. Unbiased average age-appropriate atlases for pediatric studies. NeuroImage 54, 313–327. doi:10.1016/j.neuroimage.2010.07.033

Friedman, L., Glover, G.H., 2006. Report on a multicenter fMRI quality assurance protocol. J. Magn.Reson. Imaging 23, 827–839. doi:10.1002/jmri.20583

Friedman, L., Glover, G.H., Fbirn Consortium, 2006. Reducing interscanner variability of activation in a multicenter fMRI study: controlling for signal-to-fluctuation-noise-ratio (SFNR) differences. NeuroImage 33, 471–481. doi:10.1016/j.neuroimage.2006.07.012

Giove, F., Gili, T., Iacovella, V., Macaluso, E., Maraviglia, B., 2009. Images-based suppression of unwanted global signals in resting-state functional connectivity studies. Magnetic Resonance Imaging 27, 1058–1064. doi:10.1016/j.mri.2009.06.004

Gour, N., Ranjeva, J.-P., Ceccaldi, M., Confort-Gouny, S., Barbeau, E., Soulier, E., Guye, M., Didic, M., Felician, O., 2011. Basal functional connectivity within the anterior temporal network is associated with performance on declarative memory tasks. NeuroImage 58, 687–697. doi:10.1016/j.neuroimage.2011.05.090

Greicius, M.D., Srivastava, G., Reiss, A.L., Menon, V., 2004. Default-mode network activity distinguishes Alzheimer’s disease from healthy aging: evidence from functional MRI. Proc. Natl. Acad. Sci. U.S.A. 101, 4637–4642. doi:10.1073/pnas.0308627101

Guillozet, A.L., Weintraub, S., Mash, D.C., Mesulam, M.M., 2003. Neurofibrillary tangles, amyloid, and memory in aging and mild cognitive impairment. Arch Neurol 60, 729–736. doi:10.1001/archneur.60.5.729

Kelly, C., Biswal, B.B., Craddock, R.C., Castellanos, F.X., Milham, M.P., 2012. Characterizing variation in the functional connectome: promise and pitfalls. Trends in Cognitive Sciences 16, 181–188. doi:10.1016/j.tics.2012.02.001

Klunk, W.E., Price, J.C., Mathis, C.A., Tsopelas, N.D., Lopresti, B.J., Ziolko, S.K., Bi, W., Hoge, J.A.,Cohen, A.D., Ikonomovic, M.D., Saxton, J.A., Snitz, B.E., Pollen, D.A., Moonis, M., Lippa, C.F., Swearer, J.M., Johnson, K.A., Rentz, D., Fischman, A.J., Aizenstein, H.J., DeKosky, S.T., 2007. Amyloid deposition begins in the striatum of presenilin-1 mutation carriers from two unrelated pedigrees. Journal of Neuroscience 27, 6174–6184. doi:10.1523/JNEUROSCI.0730-07.2007

Koch, W., Teipel, S., Mueller, S., Benninghoff, J., Wagner, M., Bokde, A.L.W., Hampel, H., Coates, U., Reiser, M., Meindl, T., 2012. Diagnostic power of default mode network resting state fMRI in the detection of Alzheimer’s disease. Neurobiology of Aging 33, 466–478. doi:10.1016/j.neurobiolaging.2010.04.013

Liang, P., Wang, Z., Yang, Y., Li, K., 2012. Three subsystems of the inferior parietal cortex are differently affected in mild cognitive impairment. J. Alzheimers Dis. 30, 475–487. doi:10.3233/JAD-2012-111721

Lim, H.K., Nebes, R., Snitz, B., Cohen, A., Mathis, C., Price, J., Weissfeld, L., Klunk, W., Aizenstein, H.J., 2014. Regional amyloid burden and intrinsic connectivity networks in cognitively normal elderly subjects. Brain 137, 3327–3338. doi:10.1093/brain/awu271

Lund, T.E., Madsen, K.H., Sidaros, K., Luo, W.-L., Nichols, T.E., 2006. Non-white noise in fMRI: Does modelling have an impact? NeuroImage 29, 54–66. doi:10.1016/j.neuroimage.2005.07.005

Madsen, S.K., Ho, A.J., Hua, X., Saharan, P.S., Toga, A.W., Jack, C.R., Weiner, M.W., Thompson, P.M., for the Alzheimer’s Disease Neuroimaging Initiative, 2010. 3D maps localize caudate nucleus atrophy in 400 Alzheimer’s disease, mild cognitive impairment, and healthy elderly subjects. Neurobiology of Aging 31, 1312–1325. doi:10.1016/j.neurobiolaging.2010.05.002

Maruyama, M., Shimada, H., Suhara, T., Shinotoh, H., Ji, B., Maeda, J., Zhang, M.-R., Trojanowski, J.Q., Lee, V.M.Y., Ono, M., Masamoto, K., Takano, H., Sahara, N., Iwata, N., Okamura, N.,Furumoto, S., Kudo, Y., Chang, Q., Saido, T.C., Takashima, A., Lewis, J., Jang, M.-K., Aoki, I., Ito, H., Higuchi, M., 2013. Imaging of Tau Pathology in a Tauopathy Mouse Model and in Alzheimer Patients Compared to Normal Controls. Neuron 79, 1094–1108. doi:10.1016/j.neuron.2013.07.037

McDonald, C.R., McEvoy, L.K., Gharapetian, L., Fennema-Notestine, C., Hagler, D.J., Holland, D., Koyama, A., Brewer, J.B., Dale, A.M., for the Alzheimer’s Disease Neuroimaging Initiative, 2009. Regional rates of neocortical atrophy from normal aging to early Alzheimer disease. Neurology 73, 457–465. doi:10.1212/WNL.0b013e3181b16431

Nutter-Upham, K.E., Saykin, A.J., Rabin, L.A., Roth, R.M., Wishart, H.A., Pare, N., Flashman, L.A., 2008. Verbal fluency performance in amnestic MCI and older adults with cognitive complaints. Arch Clin Neuropsychol 23, 229–241. doi:10.1016/j.acn.2008.01.005

Packard, M.G., Knowlton, B.J., 2002. Learning and memory functions of the Basal Ganglia. Annu.Rev. Neurosci. 25, 563–593. doi:10.1146/annurev.neuro.25.112701.142937

Petersen, R.C., 2004. Mild cognitive impairment as a diagnostic entity. J. Intern. Med. 256, 183–194. doi:10.1111/j.1365-2796.2004.01388.x

Pievani, M., Agosta, F., Pagani, E., Canu, E., Sala, S., Absinta, M., Geroldi, C., Ganzola, R., Frisoni, G.B., Filippi, M., 2010. Assessment of white matter tract damage in mild cognitive impairment and Alzheimer’s disease. Hum. Brain Mapp. 31, 1862–1875. doi:10.1002/hbm.20978

Power, J.D., Barnes, K.A., Snyder, A.Z., Schlaggar, B.L., Petersen, S.E., 2012. Spurious but systematic correlations in functional connectivity MRI networks arise from subject motion. NeuroImage 59, 2142–2154. doi:10.1016/j.neuroimage.2011.10.018

Qi, Z., Wu, X., Wang, Z., Zhang, N., Dong, H., Yao, L., Li, K., 2010. Impairment and compensation coexist in amnestic MCI default mode network. NeuroImage 50, 48–55. doi:10.1016/j.neuroimage.2009.12.025

Rémy, F., Vayssière, N., Saint-Aubert, L., Barbeau, E., Pariente, J., 2015. White matter disruption at the prodromal stage of Alzheimer’s disease: Relationships with hippocampal atrophy and episodic memory performance. NeuroImage Clin 7, 482–492. doi:10.1016/j.nicl.2015.01.014

Saad, Z.S., Gotts, S.J., Murphy, K., Chen, G., Jo, H.J., Martin, A., Cox, R.W., 2012. Trouble at rest: how correlation patterns and group differences become distorted after global signal regression. Brain Connectivity 2, 25–32. doi:10.1089/brain.2012.0080

Shehzad, Z., Kelly, C., Reiss, P.T., Cameron Craddock, R., Emerson, J.W., McMahon, K., Copland, D.A., Castellanos, F.X., Milham, M.P., 2014. A multivariate distance-based analytic framework for connectome-wide association studies. NeuroImage 93 Pt 1, 74–94. doi:10.1016/j.neuroimage.2014.02.024

Sheline, Y.I., Morris, J.C., Snyder, A.Z., Price, J.L., Yan, Z., D’Angelo, G., Liu, C., Dixit, S., Benzinger, T., Fagan, A., Goate, A., Mintun, M.A., 2010a. APOE4 Allele Disrupts Resting State fMRI Connectivity in the Absence of Amyloid Plaques or Decreased CSF A 42. Journal of Neuroscience 30, 17035–17040. doi:10.1523/JNEUROSCI.3987-10.2010

Sheline, Y.I., Raichle, M.E., Snyder, A.Z., Morris, J.C., Head, D., Wang, S., Mintun, M.A., 2010b. Amyloid Plaques Disrupt Resting State Default Mode Network Connectivity in Cognitively Normal Elderly. Biological Psychiatry 67, 584–587. doi:10.1016/j.biopsych.2009.08.024

Small, G.W., Kepe, V., Ercoli, L.M., Siddarth, P., Bookheimer, S.Y., Miller, K.J., Lavretsky, H., Burggren, A.C., Cole, G.M., Vinters, H.V., Thompson, P.M., Huang, S.-C., Satyamurthy, N., Phelps, M.E., Barrio, J.R., 2006. PET of brain amyloid and tau in mild cognitive impairment. N. Engl. J. Med. 355, 2652–2663. doi:10.1056/NEJMoa054625

Sorg, C., Riedl, V., Mühlau, M., Calhoun, V.D., Eichele, T., Läer, L., Drzezga, A., Förstl, H., Kurz, A., Zimmer, C., Wohlschläger, A.M., 2007. Selective changes of resting-state networks in individuals at risk for Alzheimer’s disease. Proc. Natl. Acad. Sci. U.S.A. 104, 18760–18765. doi:10.1073/pnas.0708803104

Van Dijk, K.R.A., Hedden, T., Venkataraman, A., Evans, K.C., Lazar, S.W., Buckner, R.L., 2010. Intrinsic functional connectivity as a tool for human connectomics: theory, properties, and optimization. Journal of Neurophysiology 103, 297–321. doi:10.1152/jn.00783.2009

Vanhoutte, G., Verhoye, M., Van der Linden, A., 2006. Changing body temperature affects the T2* signal in the rat brain and reveals hypothalamic activity. Magn Reson Med 55, 1006–1012. doi:10.1002/mrm.20861

Villemagne, V.L., Ataka, S., Mizuno, T., Brooks, W.S., Wada, Y., Kondo, M., Jones, G., Watanabe, Y., Mulligan, R., Nakagawa, M., Miki, T., Shimada, H., O’Keefe, G.J., Masters, C.L., Mori, H., Rowe, C.C., 2009. High striatal amyloid beta-peptide deposition across different autosomal Alzheimer disease mutation types. Arch Neurol 66, 1537–1544. doi:10.1001/archneurol.2009.285

Villeneuve, S., Jagust, W.J., 2015. Imaging Vascular Disease and Amyloid in the Aging Brain: Implications for Treatment. J Prev Alzheimers Dis 2, 64–70. doi:10.14283/jpad.2015.47

Willer, C.J., Li, Y., Abecasis, G.R., 2010. METAL: fast and efficient meta-analysis of genomewide association scans. Bioinformatics 26, 2190–2191. doi:10.1093/bioinformatics/btq340

Wu, L., Soder, R.B., Schoemaker, D., Carbonnell, F., Sziklas, V., Rowley, J., Mohades, S., Fonov, V., Bellec, P., Dagher, A., Shmuel, A., Jia, J., Gauthier, S., Rosa-Neto, P., 2014. Resting state executive control network adaptations in amnestic mild cognitive impairment. J. Alzheimers Dis. 40, 993–1004. doi:10.3233/JAD-131574

Yan, C.-G., Liu, D., He, Y., Zou, Q., Zhu, C., Zuo, X., Long, X., Zang, Y., 2009. Spontaneous Brain Activity in the Default Mode Network Is Sensitive to Different Resting-State Conditions with Limited Cognitive Load. PLoS ONE 4, e5743. doi:10.1371/journal.pone.0005743

Zhang, H.-Y., Wang, S.-J., Liu, B., Ma, Z.-L., Yang, M., Zhang, Z., Teng, G.-J., 2010. Resting brain connectivity: changes during the progress of Alzheimer disease. Radiology 256, 598–606. doi:10.1148/radiol.10091701

